# Reference-Gene Stability for RT-qPCR in the Medial Prefrontal Cortex and Hippocampal Subregions of the 5xFAD Mouse Brain

**DOI:** 10.64898/2026.10.07.756185

**Authors:** A.R. Ilina, V.A. Nikitina, A.N. Trofimov, I.N. Abdurasulova, G.A. Ryzhak, O.V. Shamova

## Abstract

Reverse transcription quantitative real-time PCR (RT-qPCR) requires normalization of the data to stably expressed reference genes, whose suitability must be validated for each experimental model and tissue. The present work aimed to assess the expression stability of seven candidate reference genes (*Ppia, Ywhaz, Rpl27, Pgk1, Sdha, Gapdh, Actb*) in three brain structures, the medial prefrontal cortex, the dorsal and ventral hippocampus, in 5xFAD mice, a model of Alzheimer’s disease, and their wild-type controls. In addition to the selection of stable reference genes, we also propose combining reference genes into multiplex assays, which reduces the number of individual reactions while preserving normalization accuracy. mRNA levels were measured by RT-qPCR using hydrolysis probes, and expression stability was analyzed separately for each brain structure using the online tool RefFinder, which combines four algorithms (comparative Delta-Ct, BestKeeper, NormFinder and geNorm) to compute a comprehensive ranking.

Reference gene stability proved to be region-specific. The most stable pairs were *Ywhaz* and *Actb* for the medial prefrontal cortex, *Gapdh* and *Actb* for the dorsal hippocampus, and *Rpl27* and *Pgk1* for the ventral hippocampus. *Actb* ranked among the most stable genes in the cortex and dorsal hippocampus but among the least stable in the ventral hippocampus, whereas *Ppia* was the least stable throughout. Expression of most candidate genes did not differ between wild-type and 5xFAD mice. The exception was *Ywhaz*, which was reduced in the ventral hippocampus of 5xFAD mice, consistent with its low stability ranking there. These results demonstrate the region-specific nature of reference gene selection in the 5xFAD mouse and allow us to recommend specific gene sets for RT-qPCR normalization in the medial prefrontal cortex and the dorsal and ventral hippocampus.

## INTRODUCTION

Reverse transcription quantitative real-time PCR (RT-qPCR) remains one of the most widely used and sensitive methods for assessing gene expression at the mRNA level. However, this sensitivity is also a limitation, since the results can be substantially distorted by variation between samples in the quantity and quality of the starting RNA, in reverse transcription efficiency, and in reaction setup. To remove these technical differences, data are normalized to endogenous reference genes, whose expression should remain stable regardless of the experimental treatment and the tissue under study [1, 2]. If a gene that is unstable under the given conditions is chosen for normalization, small but biologically meaningful changes in target gene expression may be missed or, conversely, falsely inflated [3, 4].

Despite this, “classical” housekeeping genes such as GAPDH and β-actin (Actb) are often used as reference genes by default, without prior verification of their stability. Yet a considerable body of evidence indicates that the expression of these genes varies with species, tissue type, developmental stage and experimental treatment, which makes their blind selection unreliable [5, 6, 7]. It is therefore recommended that, for each experimental model, the stability of several candidate genes be assessed beforehand and that normalization be performed to the geometric mean of the two or more most stable genes [3]. The need for such validation has repeatedly been confirmed for brain tissue as well, where reference genes are selected separately for the specific conditions of a given study [8, 9]. A similar need has been demonstrated for human neural cells derived from induced pluripotent stem cells: none of the 16 candidate genes proved universally suitable across all stages of neural differentiation, so that a distinct set of genes was recommended for normalization at each stage [10].

The correct choice of reference genes becomes particularly important when studying the brain under neurodegenerative conditions. Alzheimer’s disease (AD), the most common neurodegenerative disorder, is accompanied by β-amyloid accumulation, neuroinflammation and reactive gliosis, which may affect the expression of a wide range of genes, potentially including housekeeping genes [11, 12]. Transgenic 5xFAD mice reproduce key features of early amyloid pathology and are widely used to study the molecular mechanisms of the disease [13, 14, 15]. Prior validation of candidate reference genes is therefore required before analyzing target gene expression in this model.

A further source of variability is the regional heterogeneity of the brain. In particular, the dorsal and ventral hippocampus differ in their transcriptomic and functional properties [16], so that the stability of the same reference gene may differ between individual brain structures. This means that a set of reference genes intended to be universal across the whole brain may prove inapplicable, and that the selection should be carried out separately for each structure under study [17, 18, 19].

Although reference genes have been validated for the human brain in neurodegenerative disease [8] and in individual structures of the mouse brain [9, 18], such validation had until recently not been performed for the 5xFAD model of Alzheimer’s disease. It was recently carried out by Daini et al. [20], who assessed the stability of reference genes in the cortex and hippocampus of 5xFAD mice with respect to age and sex. Nevertheless, no set of reference genes validated separately for the medial prefrontal cortex and, in particular, for the dorsal and ventral hippocampus – as transcriptomically and functionally distinct structures – has previously been proposed for 5xFAD mice. Moreover, the use of hydrolysis probes carrying different fluorophores makes it possible to combine several reference genes in a single reaction (multiplexing), reducing the number of individual qPCR assays while preserving normalization accuracy; this format has so far rarely been applied to reference gene validation in the brain.

In the present work, we assessed the expression stability of seven candidate reference genes (*Ppia, Ywhaz, Rpl27, Pgk1, Sdha, Gapdh, Actb*) in three brain structures – the medial prefrontal cortex and the dorsal and ventral hippocampus – in 5xFAD mice and their wild-type controls, and to select the most stable genes for RT-qPCR normalization in each of these structures, including their combination into multiplex panels.

## METHODS

### Animals

The study was carried out using the transgenic 5xFAD mouse line, whose genome expresses a triple-mutant human APP and a double-mutant human PSEN1, carrying five mutations in total: three in the APP gene (K670N/M671L, Swedish; I716V, Florida; V717I, London) and two in the PSEN1 gene (M146L and L286V), which are associated with the development of familial AD. The 5xFAD model was obtained by the Saint-Petersburg Institute of Bioregulation and Gerontology from the Centre for Preclinical Testing of the Institute of Physiologically Active Substances RAS (Chernogolovka, Moscow region, Russia). The 5xFAD transgenes were maintained on a C57BL/6 × SJL background by crossing with female C57BL/6 × SJL (B6SJL) mice received by the Saint-Petersburg Institute of Bioregulation and Gerontology from the Nursery for laboratory animals: Shemyakin-Ovchinnikov Institute of bioorganic chemistry (Pushchino, Moscow Region, Russia).

Animals were housed under a controlled microclimate (temperature 22 ± 2 °C, relative humidity 50–60%, 12/12 h light/dark cycle), with access to water and standard pelleted chow ad libitum in the vivarium of the Institute of Experimental Medicine. All procedures involving animals were performed in accordance with the European Convention for the Protection of Vertebrate Animals used for Experimental and Other Scientific Purposes (Strasbourg, 1986).

The study was performed on brain samples from male wild-type mice (WT, n = 7) and transgenic 5xFAD mice (5xFAD, n = 7). The genotype of the offspring was determined by genotyping; transgene-negative littermates were used as wild-type controls (WT). Three structures were dissected from each brain: the medial prefrontal cortex (mPFC) and the dorsal (DH) and ventral (VH) hippocampus.

### Genotyping

Genotyping was performed on genomic DNA extracted from mouse tail tips using a genomic DNA isolation kit (DU-250, Biolabmix, Russia). PCR was carried out in a 2.5× reaction mix containing SYBR Green I dye (Syntol, Russia) and a set of specific oligonucleotide primers (olMR1644, 5′-AATAGAGAACGGCAGGAGCA-3′; olMR1645, 5′-GCCATGAGGGCACTAATCAT-3′; olMR3610, 5′-AGGACTGACCACTCGACCAG-3′; olMR3611, 5′-CGGGGGTCTAGTTCTGCAT-3′; GPDH-F, 5′-CCACTCAGAAGACTGTGGAT-3′ and GPDH-R, 5′-GTCATCATACTTGGCAGGTT-3′ as an internal control) under the following amplification protocol: 50 °C for 10 min and 95 °C for 3 min, followed by 45 cycles of 95 °C for 10 s, 58 °C for 40 s and 72 °C for 1 s (+ plate read), with melting-curve analysis from 50 to 95 °C (0.5 °C increments, 5 s each).

### Brain structure dissection

Brain structures were dissected from sections prepared on a Thermo Scientific Microm HM 525 cryostat (Thermo Fisher Scientific, USA) at −20 °C, in accordance with the atlas of Paxinos and Franklin (2019). The dissected structures were frozen and stored at −70 °C until RNA isolation.

### RNA isolation

Total RNA was isolated by the single-step acid guanidinium thiocyanate–phenol–chloroform extraction method [22]. Tissue was homogenized with polypropylene pestles on ice using ExtractRNA reagent (Evrogen, Moscow, Russia) according to the manufacturer’s instructions. The homogenate was incubated for 5 min at room temperature and centrifuged for 10 min (+4 °C, 12 000 rpm); chloroform (1/5 volume of ExtractRNA) was added to the supernatant, which was mixed, incubated for 20 min at room temperature and centrifuged for 20 min (+4 °C, 12 000 rpm) to separate the phases. The upper aqueous phase was mixed with isopropanol (1/2 volume of ExtractRNA), and RNA was precipitated overnight at −20 °C. RNA was pelleted by centrifugation (30 min, +4 °C, 12 000 rpm), washed with 75% ethanol and stored in 75% ethanol at −20 °C. RNA concentration and purity were assessed on a NanoDrop 2000 spectrophotometer (Thermo Fisher Scientific, USA) by absorbance at 260 nm and the A260/280 and A260/230 ratios; only samples with an A260/280 ratio of 1.8–2.1 were used for further analysis.

### Reverse transcription

First-strand cDNA was synthesized from 1 ng of total RNA with the MMLV RT kit (Evrogen, Moscow, Russia; Cat. No. SK021), following the manufacturer’s instructions. RNA was first denatured at 60 °C for 3 min. The reverse-transcription reaction (20 µL total) then combined 4 µL of 5× first-strand buffer, 2 µL of 10 mM dNTP mix, 1 µL of random (dN)_10_ primer, 2 µL of DTT and 0.5 µL of MMLV reverse transcriptase. Synthesis proceeded at 37 °C for 60 min, after which the enzyme was inactivated at 70 °C for 2 min. The resulting cDNA was kept at −20 °C until analysis.

### Real-time PCR

The reaction was carried out in a final volume of 10 µl containing MgCl_2_ (25 mM), dNTP mix (10 mM each; Evrogen), Taq DNA polymerase (Alkor Bio), forward and reverse primers, fluorescent probes (Alkor Bio), 10× Taq polymerase buffer (Alkor Bio), 1 µl of cDNA and dH_2_O. Detection was performed with hydrolysis (TaqMan) probes carrying a FAM, HEX or ROX fluorescent reporter at the 5′ end and a BHQ-1/BHQ-2 quencher at the 3′ end, which allowed several targets to be detected in a single reaction (multiplexing). The primer and probe sequences, reporters and quenchers for each reference gene are given in Table S1. The final concentrations of primers, probes, enzyme and MgCl_2_ were optimized for each multiplex. Amplification was performed on a C1000 Touch Thermal Cycler with a CFX96 Touch Real-Time PCR Detection System (Bio-Rad Laboratories, USA), and the data were processed in CFX Manager 2.1 using the threshold method. All samples were analysed in duplicate, and Cq values were averaged.

The amplification programme was as follows: enzyme activation at 94 °C for 180 s, followed by 4 cycles (94 °C for 10 s; 60 °C for 20 s; 72 °C for 5 s) and 44 cycles (94 °C for 10 s; 60 °C for 20 s with fluorescence acquisition; 72 °C for 5 s).

Primers and probes were designed in silico (IDT OligoAnalyzer, Primer-BLAST, Primer3Plus and Oligo 7; reference sequences from NCBI RefSeq and the UCSC Genome Browser) within conserved regions spanning an exon–exon junction by at least 3 nucleotides. The selection criteria were: GC content 40–60%; primer length 16–26 bp and probe length 18–28 bp; primer melting temperature 63–66 °C and probe melting temperature 68–72 °C; the 5′-terminal nucleotide of the probe not a guanine; and thermodynamic calculations based on the nearest-neighbour model. The specificity of the designed assays was confirmed using NCBI BLAST.

### Calibrators and amplification efficiency

All samples from the three brain structures under study were analysed within a single PCR run on one plate, which minimized inter-run variability. Real-time PCR was performed in a multiplex format, with the candidate reference genes grouped into separate panels (Sdha, Pgk1, Gapdh; Actb; Ywhaz, Rpl27; Ppia). Amplification efficiency (E) was determined from a standard curve constructed on a series of two-fold dilutions of calibrator cDNA. E was calculated from the slope of the relationship between the replicate-averaged Cq and log_10_(relative quantity) using the formula E = 10^(−1/slope)^ − 1 . The slope and the coefficient of determination (R^2^) of the standard curve were calculated from the averaged Cq values of each dilution. The E and R^2^ values are given in Fig. S4.

### Reference gene stability analysis

The expression stability of the seven candidate reference genes (*Ppia, Ywhaz, Rpl27, Pgk1, Sdha, Gapdh, Actb*), selected on the basis of published data on the mouse brain, was assessed separately for each structure using the online tool RefFinder. RefFinder combines four algorithms — the comparative Delta-Ct method, BestKeeper, NormFinder and geNorm — and computes a comprehensive ranking as the geometric mean of the ranks obtained by each method. Samples of both genotypes were analyzed together within each structure so that expression stability would also reflect robustness to genotype differences. For normalization within each structure, the geometric mean of the two most stable reference genes was used, in accordance with the recommendation of [3].

### Quality control and exclusion of technical outliers

Before analysis, the median Cq across all reference genes was calculated for each sample; samples deviating from the group median by more than three median absolute deviations were excluded as technically inadequate [23]. By this criterion, two samples were excluded — one in the cortex (WT) and one in the ventral hippocampus (5xFAD) — in which the Cq values of all seven genes were uniformly elevated, indicating insufficient cDNA rather than biological variability. The final sample numbers were: mPFC, 13; DH, 14; VH, 13.

Data analysis and statistics Relative expression was calculated by the 2^−ΔΔCq^ method [24]. For each sample, ΔCq was computed as ΔCq = Cq(gene) − (Cq_ref1_ + Cq_ref2_)/2. Normalization was performed to the geometric mean of the relative quantities of the two most stable reference genes of the corresponding structure [3], which is equivalent to the arithmetic mean of their Cq values. ΔΔCq was then calculated as ΔΔCq = ΔCq(sample) − mean ΔCq of the WT group (calibrator), and the relative mRNA quantity was obtained as 2^−ΔΔCq^. Amplification efficiencies (Fig. S4) are reported as a quality control of the assays and were not used in the relative expression calculation.

Before comparing groups, the relative expression values of each gene in each structure were checked for outliers by the ROUT method (Q = 5%) in GraphPad Prism 8.0.1, separately for the WT and 5xFAD groups. Differences in expression between the WT and 5xFAD groups were assessed by the nonparametric Mann–Whitney U test, given the small group sizes. Differences were considered significant at *p* < 0.05. Statistical analysis and graph plotting were performed in Microsoft Excel and GraphPad Prism 8.0.1.

## RESULTS

### Reference gene stability ranking across mouse brain structures

The expression stability of the seven candidate reference genes (*Ppia, Ywhaz, Rpl27, Pgk1, Sdha, Gapdh, Actb*) was analyzed in three brain structures — the medial prefrontal cortex and the dorsal and ventral hippocampus — in 5xFAD and wild-type (B6SJL) mice using the online tool RefFinder, which combines four algorithms (comparative Delta-Ct, BestKeeper, NormFinder and geNorm). The stability values obtained by each method are shown in Fig. 1, and the final rankings in Fig. 2. The distribution of stability values across structures was consistent regardless of the algorithm (Fig. 1). Overall, the genes examined showed the most stable expression in the medial prefrontal cortex and the least stable in the dorsal hippocampus. Per-structure stability rankings by all four algorithms are shown in Figs S1–S3 for the medial prefrontal cortex, dorsal hippocampus and ventral hippocampus, respectively.

**Fig. 1.**
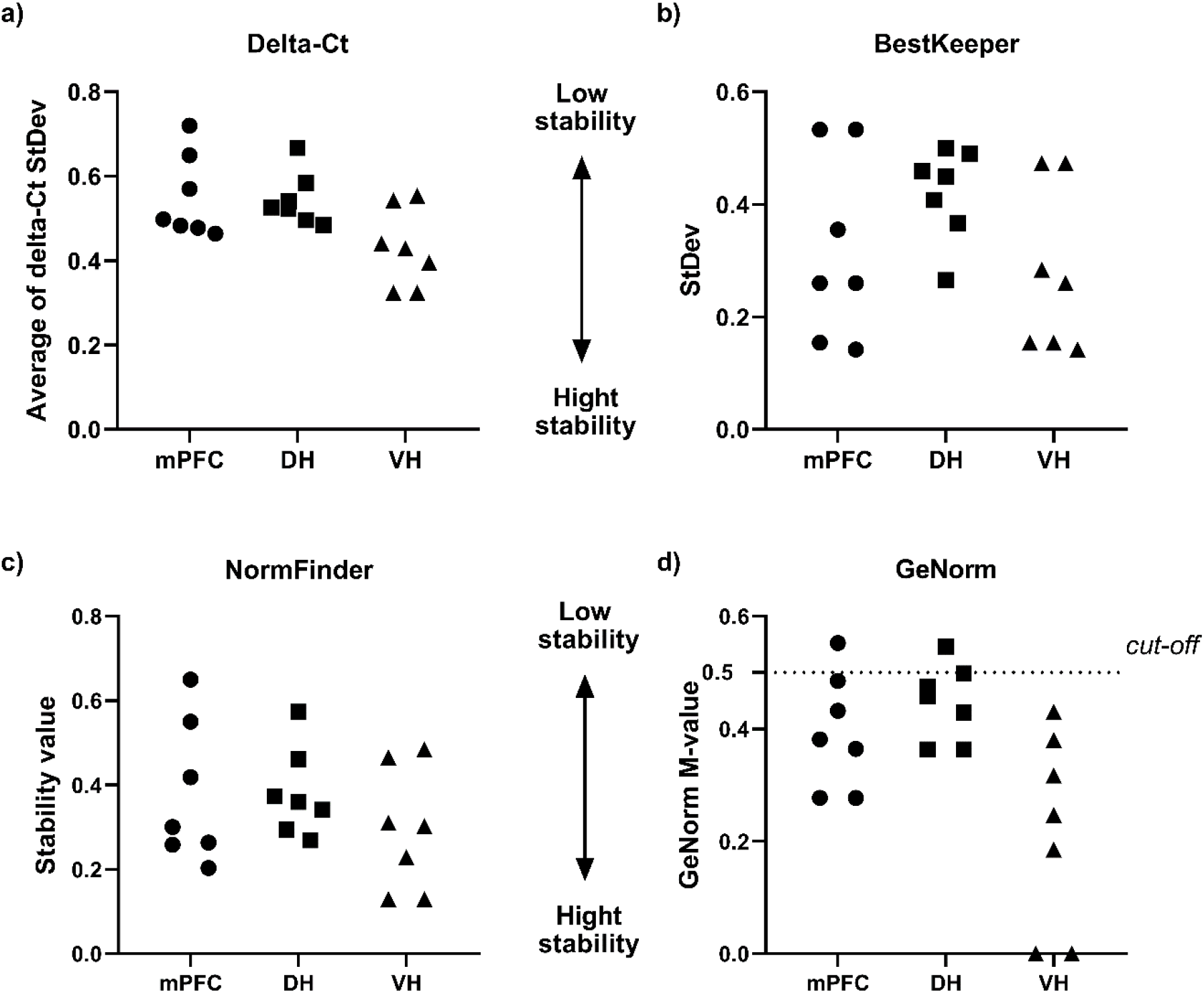
Reference gene expression stability values obtained by four methods in three brain structures of 5xFAD and wild-type mice. The expression stability values of the seven candidate genes (*Ppia, Ywhaz, Rpl27, Pgk1, Sdha, Gapdh, Actb*; each gene is represented by one point per structure) in the medial prefrontal cortex (mPFC, circles), the dorsal (DH, squares) and ventral (VH, triangles) hippocampus were analysed by (a) the comparative Delta-Ct method, (b) BestKeeper, (c) NormFinder and (d) geNorm. For geNorm (d), values above the M-value threshold of 0.5 (dashed line, cut-off) correspond to unstably expressed genes unsuitable for RT-qPCR data normalization

**Fig. 2.**
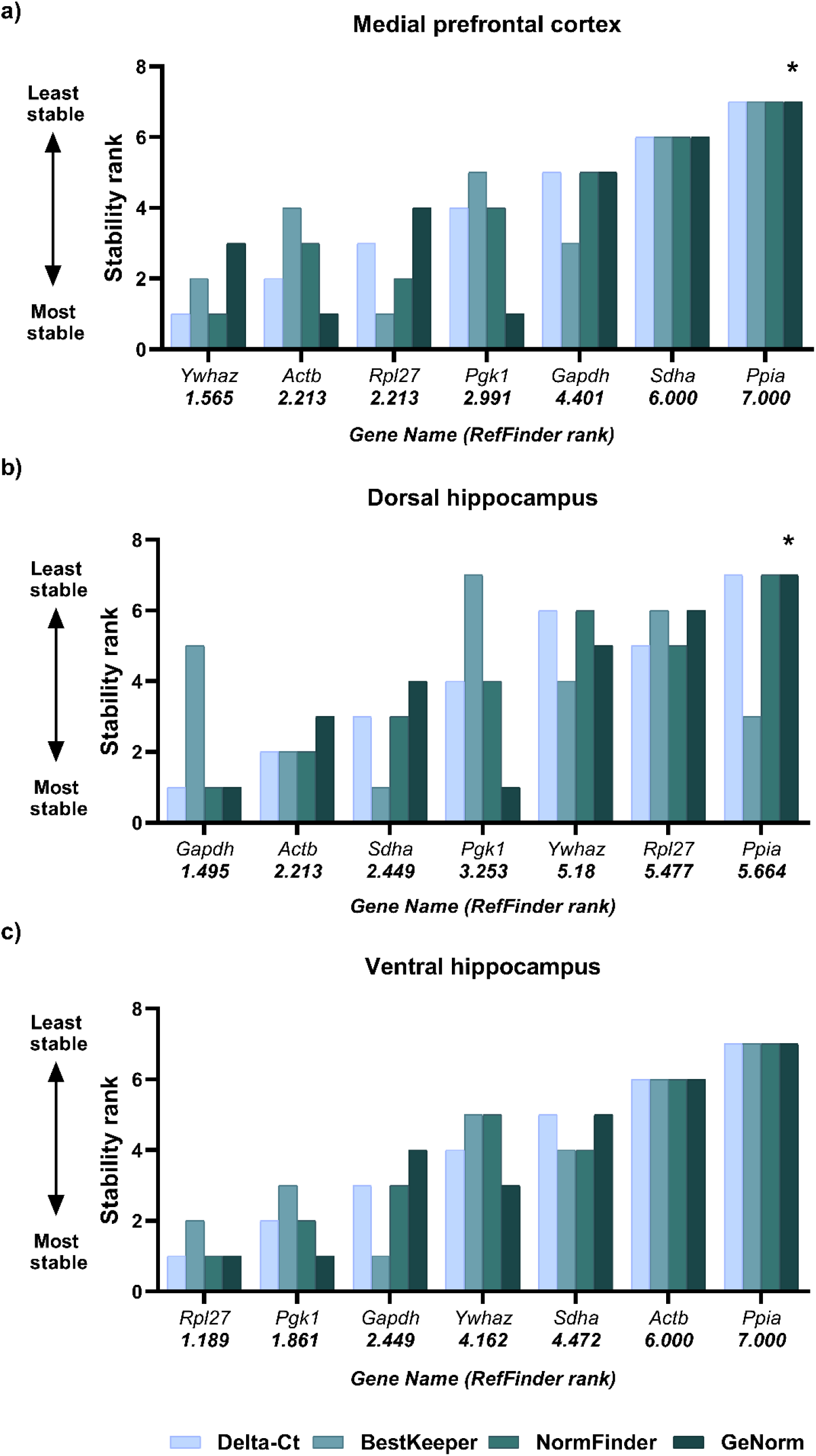
Ranking of reference gene expression stability in three brain structures of 5xFAD and wild-type mice. The expression stability of the reference genes in (a) the medial prefrontal cortex, (b) the dorsal and (c) the ventral hippocampus was analyzed using the online tool RefFinder. The comprehensive RefFinder ranking (values given in parentheses below the gene name on the x-axis) is based on the geometric mean of the ranks obtained by the four methods: comparative Delta-Ct, BestKeeper, NormFinder and geNorm. Asterisks indicate unstably expressed genes unsuitable for normalization according to the geNorm M-value threshold of 0.5.

#### Medial prefrontal cortex

According to the comprehensive RefFinder ranking, the most stable reference genes in the medial prefrontal cortex were *Ywhaz, Actb* and *Rpl27*, whereas *Ppia* showed the least stable expression (Fig. 2a). According to geNorm, *Ppia* was the only gene with an M-value above the threshold of 0.5 and was therefore unsuitable for normalization. NormFinder and the comparative Delta-Ct method likewise identified *Ywhaz* and *Actb* as the most stable, and *Ppia* and *Sdha* as the least stable. BestKeeper identified *Actb, Ywhaz* and *Rpl27* as the most stably expressed genes.

#### Dorsal hippocampus

The comprehensive RefFinder ranking identified *Gapdh, Actb* and *Sdha* as the most stable reference genes in the dorsal hippocampus, whereas *Ppia* showed the least stable expression (Fig. 2b). According to geNorm, *Ppia* had an M-value above the threshold of 0.5 and was classified as unstable and unsuitable for normalization. NormFinder and the comparative Delta-Ct method ranked *Gapdh* and *Actb* among the most stable, whereas *Ywhaz* and *Ppia* showed low stability. BestKeeper identified *Actb* and *Sdha* as the most stable.

#### Ventral hippocampus

In the ventral hippocampus, *Rpl27, Pgk1* and *Gapdh* were the most stably expressed genes, whereas *Actb* and *Ppia* were the least stable, according to the comprehensive RefFinder ranking (Fig. 2c). NormFinder and the comparative Delta-Ct method identified *Rpl27* and *Pgk1* as the most stable, and *Actb* and *Ppia* as the least stable. BestKeeper identified *Rpl27* and *Pgk1* as the most stably expressed genes. Notably, *Actb*, which ranked among the most stable genes in the medial prefrontal cortex, was among the least stable in the ventral hippocampus.

### Relative mRNA expression after normalization to optimal reference genes

The effect of genotype on the expression of the genes analyzed was assessed after normalization to the two most stable reference genes of the corresponding structure. In the medial prefrontal cortex, expression was normalized to the geometric mean of *Ywhaz* and *Actb* (Fig. 3); in the dorsal hippocampus, to *Gapdh* and *Actb* (Fig. 4); and in the ventral hippocampus, to *Rpl27* and *Pgk1* (Fig. 5). With a single exception, no statistically significant differences in relative mRNA expression were found between 5xFAD and wild-type mice (Mann–Whitney U test, *p* > 0.05), and expression levels overlapped between the groups. The exception was *Ywhaz* expression in the ventral hippocampus, which was significantly reduced in 5xFAD mice compared with wild-type (Mann–Whitney U test, *p* = 0.037; Fig. 5b). Notably, *Ywhaz* was not among the most stable genes in the ventral hippocampus and was not used for normalization in this structure; the genotype-dependence of its expression is consistent with its low stability ranking and underscores the need for region-specific selection of reference genes.

**Fig. 3.**
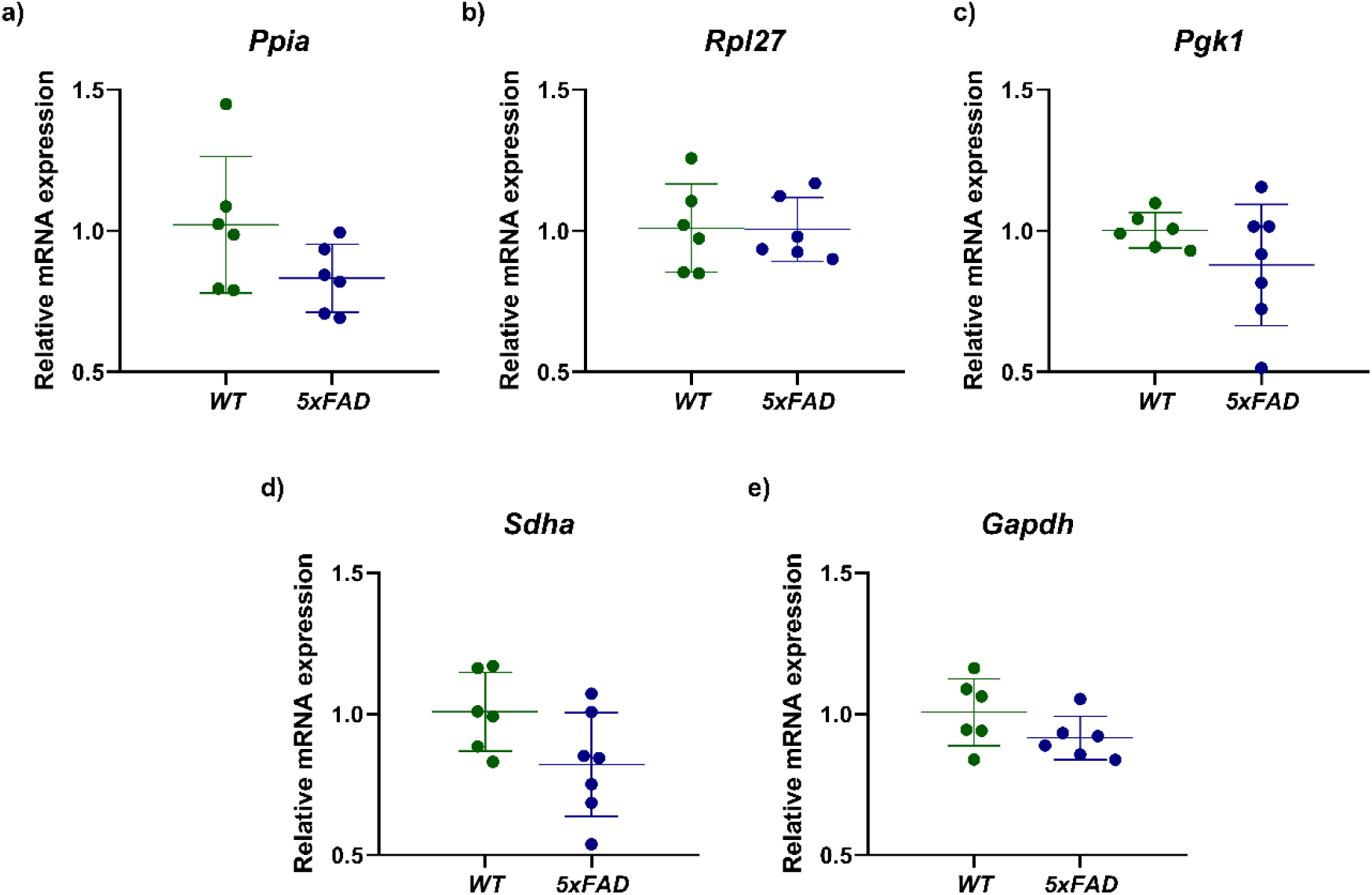
Relative mRNA expression of reference genes in the medial prefrontal cortex of 5xFAD and wild-type mice. The relative mRNA expression of (a) *Ppia*, (b) *Rpl27*, (c) *Pgk1*, (d) *Sdha* and (e) *Gapdh* in the medial prefrontal cortex of wild-type (WT) and 5xFAD mice was measured by RT-qPCR. Expression was normalized to the geometric mean of the two most stably expressed genes (*Ywhaz* and *Actb*). No differences between genotypes were found (Mann–Whitney U test, *p* > 0.05).

**Fig. 4.**
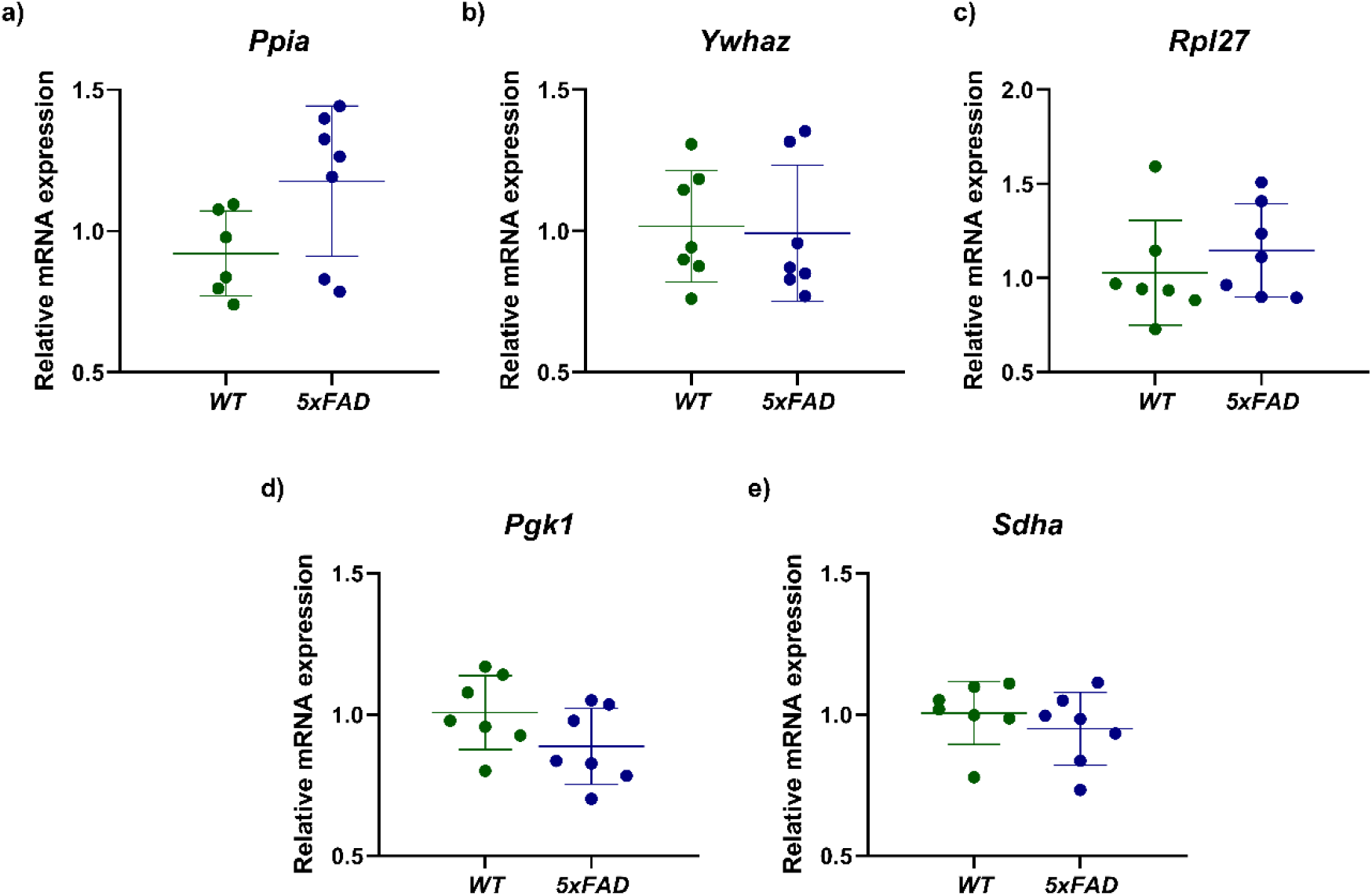
Relative mRNA expression of reference genes in the dorsal hippocampus of 5xFAD and wild-type mice. The relative mRNA expression of (a) *Ppia*, (b) *Ywhaz*, (c) *Rpl27*, (d) *Pgk1* and (e) *Sdha* in the dorsal hippocampus of wild-type (WT) and 5xFAD mice was measured by RT-qPCR. Expression was normalized to the geometric mean of the two most stably expressed genes (*Gapdh* and *Actb*). No differences between genotypes were found (Mann–Whitney U test, *p* > 0.05).

**Fig. 5.**
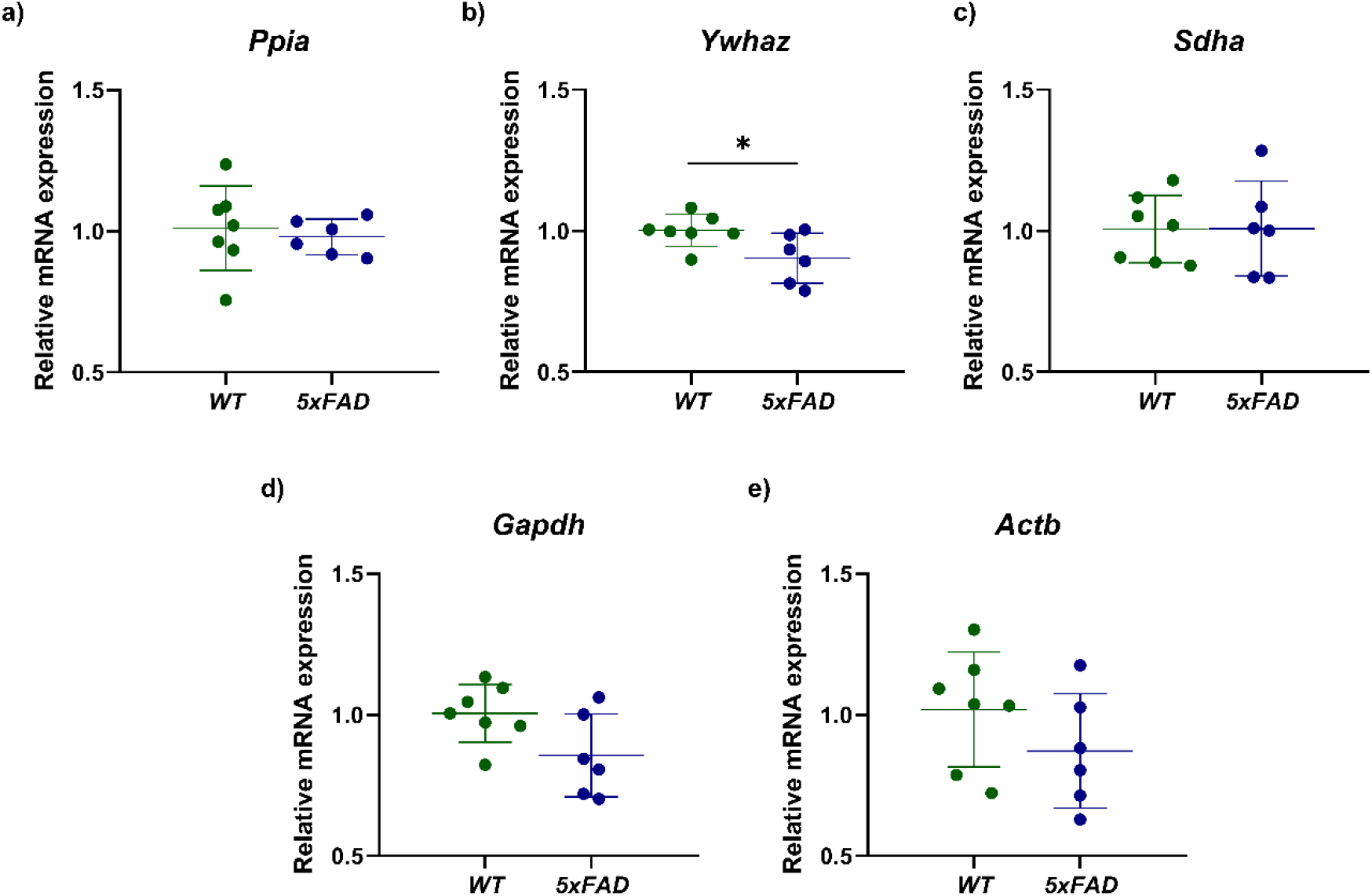
Relative mRNA expression of reference genes in the ventral hippocampus of 5xFAD and wild-type mice. The relative mRNA expression of (a) *Ppia*, (b) *Ywhaz*, (c) *Sdha*, (d) *Gapdh* and (e) *Actb* in the ventral hippocampus of wild-type (WT) and 5xFAD mice was measured by RT-qPCR. Expression was normalized to the geometric mean of the two most stably expressed genes (*Rpl27* and *Pgk1*). No statistically significant differences between genotypes were found (Mann–Whitney U test, *p* > 0.05), except for *Ywhaz* (b), whose expression was reduced in 5xFAD mice (*p* = 0.037).

Thus, the expression of the recommended reference genes was independent of genotype, confirming their suitability for RT-qPCR normalization in brain structures of 5xFAD and wild-type mice. The only gene with genotype-dependent expression was *Ywhaz* in the ventral hippocampus, which was therefore not selected as a reference gene for this structure.

## DISCUSSION

The present work shows that the expression stability of candidate reference genes in the brain of 5xFAD mice differs substantially between structures: no single gene and no single pair of genes proved optimal simultaneously for all three regions examined. Based on the comprehensive RefFinder ranking, the pairs *Ywhaz* and *Actb* for the medial prefrontal cortex, *Gapdh* and *Actb* for the dorsal hippocampus, and *Rpl27* and *Pgk1* for the ventral hippocampus are recommended for RT-qPCR normalization. This result is consistent with the widely accepted principle that the suitability of reference genes must be confirmed separately for each experimental system and tissue [1, 3], and extends it to the level of individual brain structures.

The behavior of the “classical” housekeeping genes *Actb* and *Gapdh*, which are often used for normalization without prior verification of their stability, proved particularly telling. In our study, *Actb* was among the most stable genes in the cortex and dorsal hippocampus but one of the least stable in the ventral hippocampus; *Gapdh*, by contrast, was of low stability in the cortex but among the most stable in the dorsal and ventral hippocampus. Both genes thus show pronounced region-dependence, which is consistent with previously published data on the unreliability of β-actin and the variability of GAPDH depending on tissue and conditions [6, 7]. Using both gene blindly as the sole reference for the whole brain could distort the results in the structure where it is unstable.

*Ppia* showed the lowest stability in all three structures and is therefore the least suitable reference gene in this model. By contrast, *Rpl27* was the most stable gene in the ventral hippocampus but ranked among the lowest in the dorsal hippocampus, which again highlights the region-specificity of stability and shows that even a ribosomal protein gene is not universally suitable. This differs from a number of studies in which ribosomal protein genes (e.g., Rpl13a) were among the most stable in the rat and human brain [8, 19], and once again points to the need for validation under specific conditions.

Our results are logically consistent with, and complement, previous work on reference gene selection in neurodegeneration. Thus, a study of reference genes in the rat brain under ketosis [19] likewise showed a dependence of stability on the hippocampal region; however, that study was performed in a different species and under different experimental conditions, and one of the housekeeping genes proved to be differentially expressed between groups. In a study of post-mortem human brain in neurodegenerative diseases [8], different reference genes were recommended using the same RefFinder tool, which is to be expected given the differences in species, tissue and candidate set. The closest study is the recent validation of reference genes in the same 5xFAD model [20], in which the analysis was structured along the axes of age and sex, and the cortex and hippocampus were treated as whole regions. A separate assessment of the cortex and the dorsal and ventral hippocampus was not performed.

The difference in optimal reference genes between the dorsal and ventral hippocampus is of particular interest. These subregions have long been described as functionally and transcriptomically distinct structures [16], and our data confirm this at the level of housekeeping gene expression. In practical terms, this means that the dorsal and ventral hippocampus should be treated as separate structures in gene expression analysis, rather than combined into a single “hippocampus”. Assessing stability at the level of the hippocampus as a whole average these subregions and may mask intra-structural heterogeneity: *Actb*, for example, is among the most stable genes in the dorsal hippocampus and among the least stable in the ventral hippocampus, and this difference would be lost if the subregions were combined.

The comparison of candidate gene expression between wild-type and 5xFAD mice revealed no significant differences in almost all cases. The only gene whose expression depended on genotype was *Ywhaz* in the ventral hippocampus, where its level was reduced in 5xFAD mice. This indicates that *Ywhaz* expression is sensitive to the pathological process and that this gene may acquire the properties of a target gene. *Ywhaz* encodes the 14-3-3ζ protein, one of the isoforms of the 14-3-3 family, which are abundantly expressed in neurons and involved in the maintenance of the cytoskeleton, synaptic function and anti-apoptotic signaling [25]. This isoform is directly implicated in the pathogenesis of Alzheimer’s disease: 14-3-3ζ binds to the tau protein and promotes its phosphorylation, is found within neurofibrillary tangles, and contributes to synaptic pathology through the accelerated degradation of synaptophysin [26]. Moreover, a reduction in YWHAZ protein levels has been shown in the hippocampus of patients with Alzheimer’s disease [27]. The reduction in *Ywhaz* mRNA that we observed specifically in the ventral — but not the dorsal — hippocampus of 5xFAD mice is consistent with these data and with the concept of regional specificity of vulnerability in Alzheimer’s disease. Notably, using the same RefFinder tool, Mengi Camur and colleagues assessed the expression of *Ywhaz* as a reference gene in the cortex, hippocampus and hypothalamus of Apoe^−/−^ mice (a model of systemic lipid metabolism disturbance) and found no significant effect of genotype [28]. The absence of changes in *Ywhaz* expression in that model contrasts with the reduction we observed in the ventral hippocampus of 5xFAD mice and is consistent with the notion that *Ywhaz* dysregulation is probably linked specifically to the amyloid- and tau-associated neurodegenerative process.

Regarding the recommended pairs, the absence of differences between genotypes indicates that the selected reference genes are not regulated by the pathological process and are therefore suitable for correct normalization when comparing genotypes. It is important to emphasize that stability ranking (which reflects the overall variability of a gene) and the comparison of means between groups are different characteristics: thus, Ppia, the least stable gene according to RefFinder, nevertheless showed no systematic shift in expression between genotypes, which is not a contradiction.

The present study has several limitations. First, the group sizes were relatively small (six to seven animals per group in each structure), and a nonparametric test was therefore used to compare genotypes. Second, amplification efficiencies were determined within multiplex reactions, which may account for the values near the lower limit of the recommended range for some genes. Third, the study was performed on males of a single age (8 months). This design was chosen deliberately: fixing age and sex makes it possible to isolate the assessment of region-specific reference gene stability from the established influence of these factors [20]. At the same time, this limits the transferability of the resulting gene sets to females and other age points, and combining the structural and the age–sex axes appear to be a natural direction for further work. Finally, a limited set of candidate reference genes was analysed, and it cannot be excluded that other genes might prove even more stable in particular structures.

Overall, our results demonstrate that the selection of reference genes for RT-qPCR in the 5xFAD mouse brain should take the specific structure into account, and provide researchers with validated reference gene sets for the medial prefrontal cortex and the dorsal and ventral hippocampus, thereby increasing the reliability of subsequent target gene expression analysis in this widely used model of Alzheimer’s disease.

## STATEMENTS AND DECLARATIONS

### Author Contributions

All authors contributed to the study conception and design. Study conception and methodology were developed by A.R. Ilina and V.A. Nikitina, with methodological guidance from A.N. Trofimov. Animal work, dissection of brain structures and RT-qPCR were performed by A.R. Ilina and V.A. Nikitina. Data processing and analysis were carried out by A.R. Ilina. The first draft of the manuscript was written by A.R. Ilina, and all authors commented on and revised the manuscript. Resources and supervision were provided by O.V. Shamova, G.A. Ryzhak and I.N. Abdurasulova. All authors read and approved the final manuscript.

### Data Availability

All data generated or analyzed during this study are included in this published article and its supplementary information files.

### Ethics approval

All procedures involving animals were performed in accordance with the European Convention for the Protection of Vertebrate Animals used for Experimental and Other Scientific Purposes (Strasbourg, 1986). The study was approved by the Ethics Committee of the Saint Petersburg Institute of Bioregulation and Gerontology (protocol No. 03, dated 29.03.23); the experimental procedures were carried out at the Institute of Experimental Medicine within the framework of a research collaboration between the two institutions.

### Consent to participate

Not applicable.

### Consent to publish

Not applicable.

