## Supplementary material for "Reference-Gene Stability for RT-qPCR in the Medial Prefrontal Cortex and Hippocampal Subregions of the 5xFAD Mouse Brain": Table S1, Figure S1, Figure S2, Figure S3, Figure S4

**SUPPLEMENTARY INFORMATION**

**Table S1. Primers and probes of the reference genes used for RT-qPCR.**

| Gene | RefSeq accession | Encoded protein | Forward primer (5′→3′) | Reverse primer (5′→3′) | Probe (5′→3′) | Reporter (5′) | Amplicon length, bp | Amplicon position (mRNA) | Multiplex group |
|---|---|---|---|---|---|---|---|---|---|
| ***Ppia*** | NM_008907.2 | Peptidylprolyl isomerase A | CCAAGACTGAATGGCTGGATG | TTAGAGCTGTCCACAGTCGG | AGCCATGGAGCGTTTTGGGTCCAGG | FAM | 146 | 394–539 | Singleplex |
| ***Ywhaz*** | NM_011740 | Tyrosine 3-monooxygenase/tryptophan 5-monooxygenase activation protein, zeta | TCTGCAACGATGTACTGTCTCT | AGGCTTTCTCTGGGGAGTTC | TGGCCGAGGTTGCTGCTGGT | FAM | 288 | 883–1170 | Ywhaz–Rpl27 |
| ***Rpl27*** | NM_011289.3 | Ribosomal protein L27 | ATCGCCAAGCGATCCAAGAT | TCAAAGCTGGGTCCCTGAAC | ACCACCTCATGCCCACAAGGTACTCT | HEX | 136 | 398–533 | Ywhaz–Rpl27 |
| ***Pgk1*** | NM_008828.3 | Phosphoglycerate kinase 1 | CTTTCGACCTCACGGTGTTG | ACGTTGAAGTCCACCCTCAT | CTGGACGTGAAGGGGAAGCGGGT | ROX | 105 | 127–231 | Sdha–Gapdh–Pgk1 |
| ***Sdha*** | NM_023281.1 | Succinate dehydrogenase complex, subunit A, flavoprotein (Fp) | GACATTCGACAGGGGAATGGT | GGACTCCTTCCGAGCTTCTG | ACCTGGTGGAGACCCTGGAGCTGCA | FAM | 115 | 1682–1796 | Sdha–Gapdh–Pgk1 |
| ***Gapdh*** | NM_008084.4 | Glyceraldehyde-3-phosphate dehydrogenase | TCTCTGCTCCTCCCTGTTCC | CCAATACGGCCAAATCCGTTC | TGCAGTGCCAGCCTCGTCCCGT | HEX | 108 | 3–110 | Sdha–Gapdh–Pgk1 |
| ***Actb*** | NM_007393.5 | Actin, beta | ACCTTCTACAATGAGCTGCG | CTGGATGGCTACGTACATGG | TCTGGGTCATCTTTTCACGGTTGGC | HEX | 100 | 374—473 | Singleplex |

**Figure S1. Reference gene expression stability in the medial prefrontal cortex of 5xFAD and wild-type mice.**

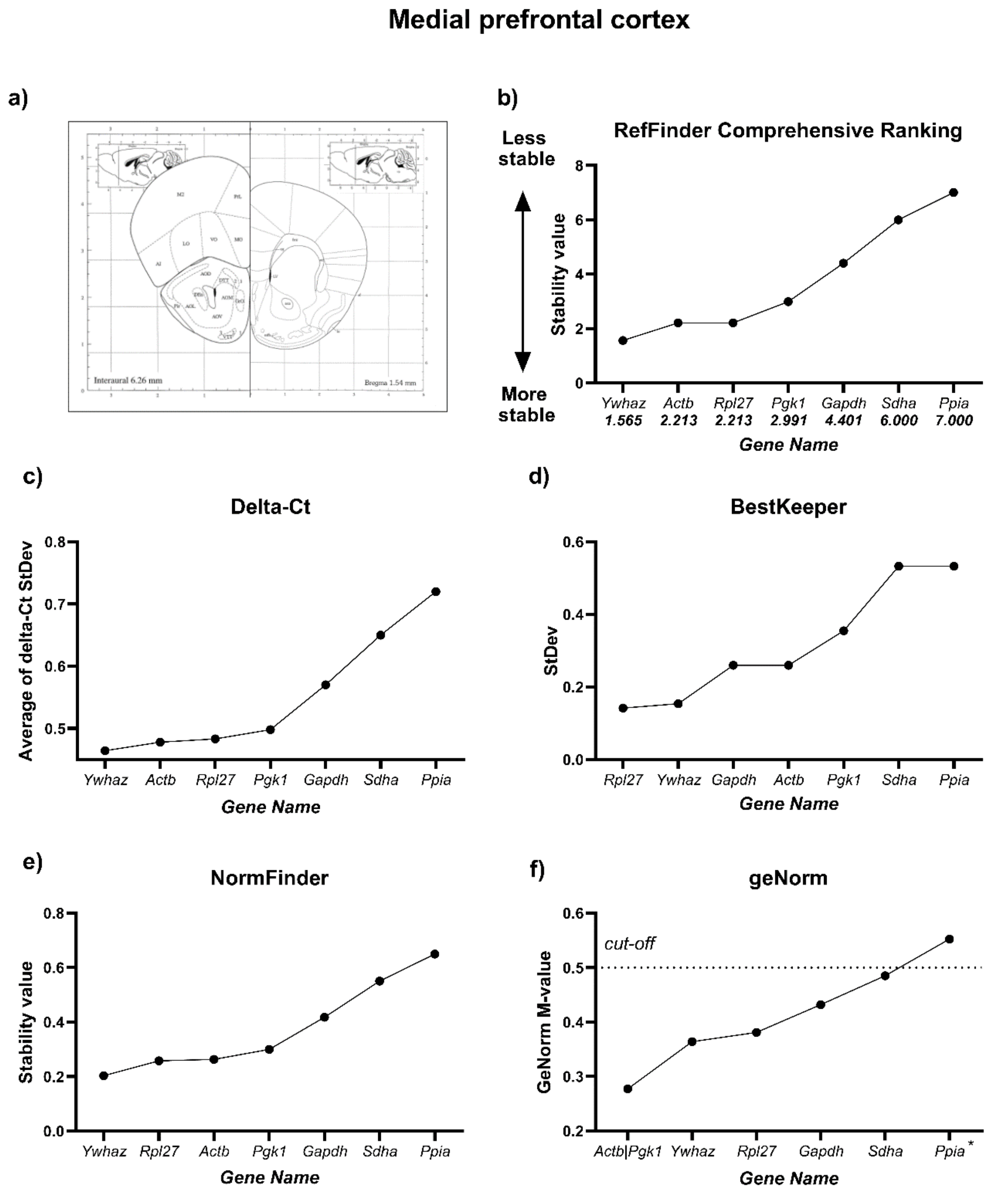

**Figure S2. Reference gene expression stability in the dorsal hippocampus of 5xFAD and wild-type mice.**

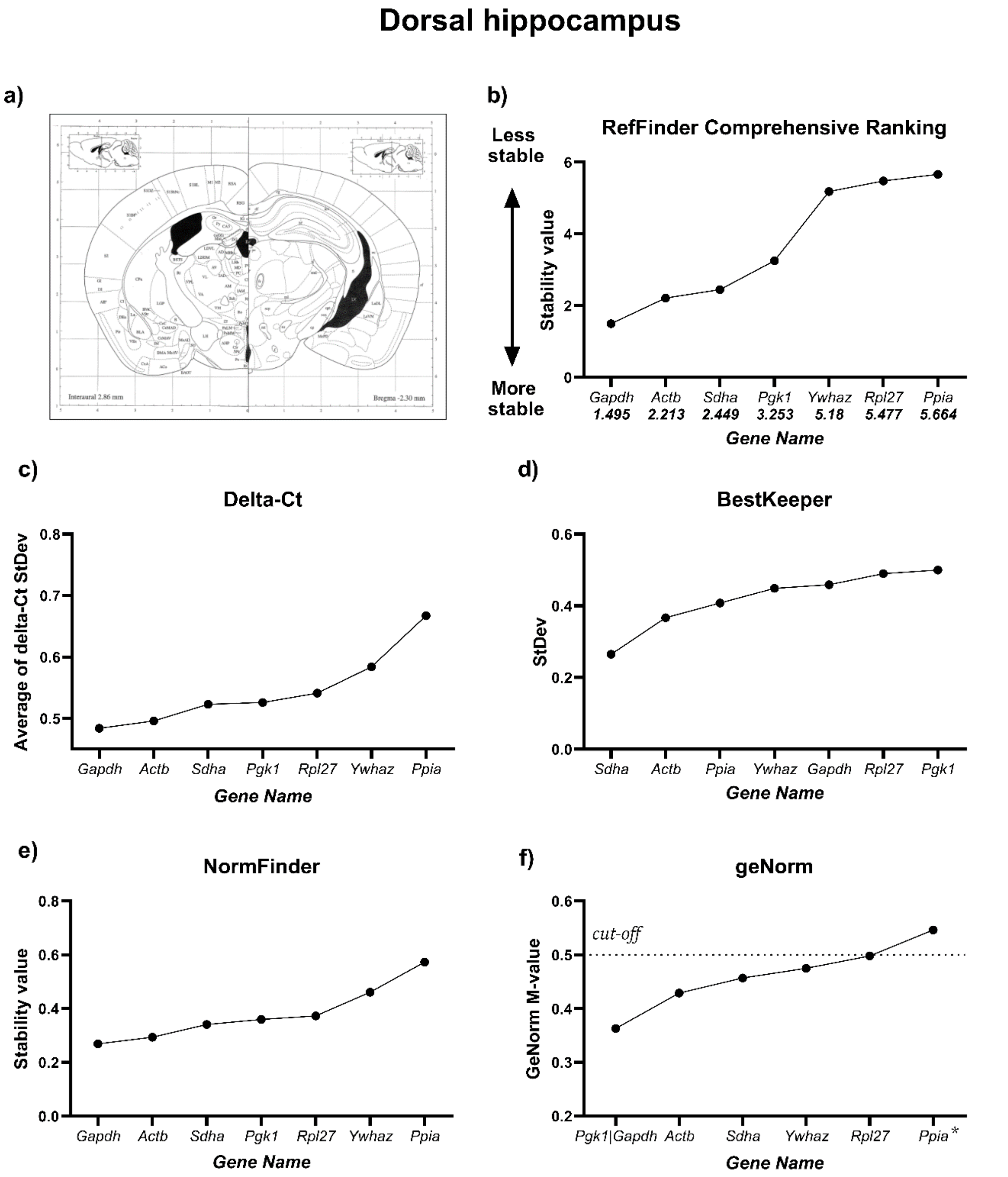

**Figure S3. Reference gene expression stability in the ventral hippocampus of 5xFAD and wild-type mice.**

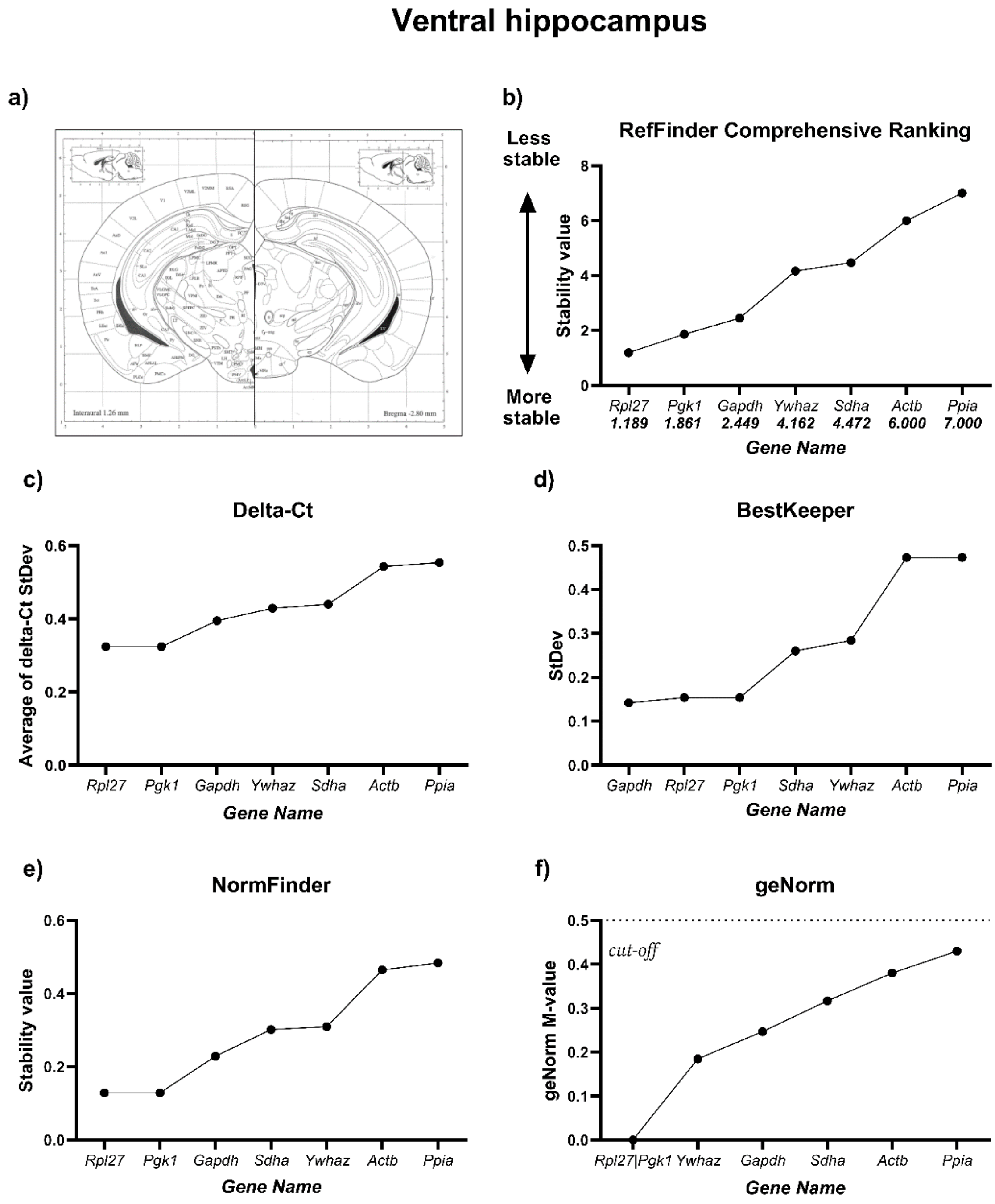

**Figure S4. Standard curves and amplification efficiencies of the reference gene assays.**

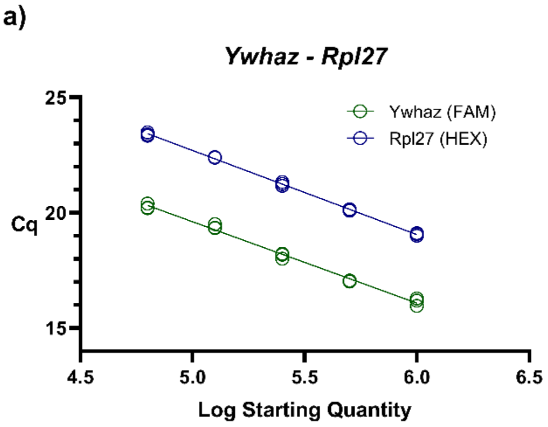


| | Slope | $R^2$ | E (%) |
|---|---|---|---|
| ***Ywhaz*** | -3.53 | 0.9966 | 91.9 |
| ***Rpl27*** | -3.66 | 0.9995 | 87.7 |

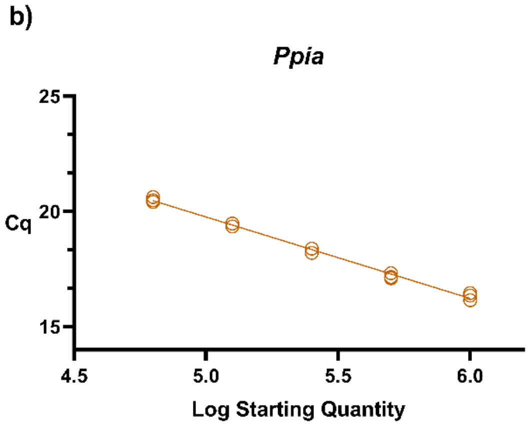


| | Slope | $R^2$ | E (%) |
|---|---|---|---|
| ***Ppia*** | -3.53 | 0.9981 | 91.93 |

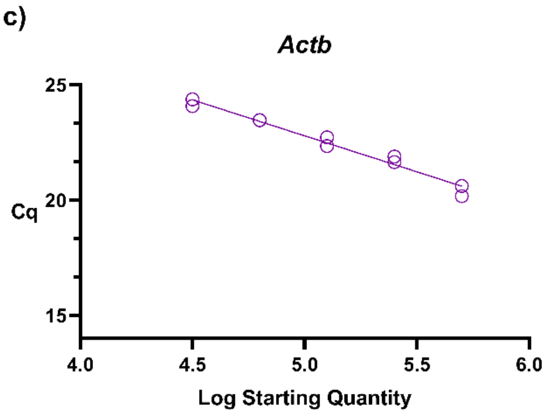


| | Slope | $R^2$ | E (%) |
|---|---|---|---|
| ***Actb*** | -3.12 | 0.9865 | 109.09 |

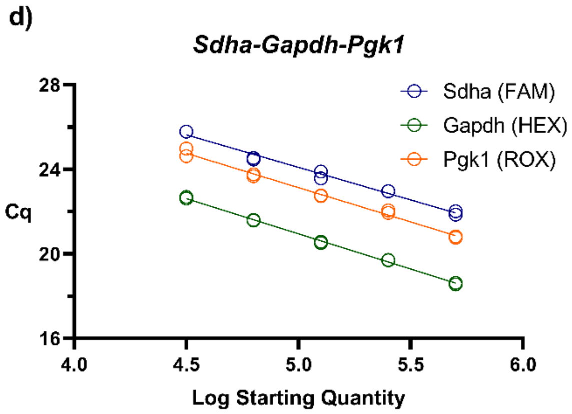


| | Slope | $R^2$ | E (%) |
|---|---|---|---|
| ***Sdha*** | -3.08 | 0.9905 | 111.02 |
| ***Gapdh*** | -3.34 | 0.9984 | 99.39 |
| ***Pgk1*** | -3.25 | 0.9959 | 103.17 |
